# Evolutionary analyses of visual opsin genes in anurans reveals diversity and positive selection suggestive of functional adaptation to distinct light environments

**DOI:** 10.1101/2021.08.27.457945

**Authors:** Ryan K Schott, Leah Perez, Matthew A Kwiatkowski, Vance Imhoff, Jennifer M Gumm

## Abstract

Among major vertebrate groups, anurans (frogs and toads) are understudied with regards to their visual systems and little is known about variation among species that differ in ecology. We sampled North American anurans representing diverse evolutionary and life histories that likely possess visual systems adapted to meet different ecological needs. Using standard molecular techniques, visual opsin genes, which encode the protein component of visual pigments, were obtained from anuran retinas. Additionally, we extracted the visual opsins from publicly available genome and transcriptome assemblies, further increasing the phylogenetic and ecological diversity of our dataset. We found that anurans consistently express four visual opsin genes (*RH1*, *LWS*, *SWS1*, and *SWS2*, but not *RH2*) even though reported photoreceptor complements vary widely among species. We found the first evidence of visual opsin duplication in an amphibian with the duplication of the *LWS* gene in the African bullfrog, which had distinct *LWS* copies on the sex chromosomes. The proteins encoded by these genes showed considerable sequence variation among species, including at sites known to shift the spectral sensitivity of visual pigments in other vertebrates and thus mediate dim-light and color vision. Using molecular evolutionary analyses of selection (d_N_/d_S_) we found significant evidence for positive selection at a subset of sites in the dim-light rod opsin gene *RH1* and the long wavelength sensitive cone opsin gene *LWS*. The function of sites inferred to be under positive selection are largely unknown, but a few are likely to affect spectral sensitivity and other visual pigment functions based on proximity to previously identified sites in other vertebrates. The observed variation cannot fully be explained by evolutionary relationships among species alone. Taken together, our results suggest that other ecological factors, such as habitat and life history, as well as behaviour, may be driving changes to anuran visual systems.

## Introduction

Anurans were used as early model systems for studies of the vertebrate visual system and many core mechanisms of visual function in vertebrates were discovered using anuran models, yet they have largely fallen out of use in vision biology (for a review see Donner and Yovanovich 2020). Relatively few modern studies have examined anuran visual systems despite the importance of vision to many aspects of anuran biology, including movement patterns, habitat preferences, foraging, reproduction, and possibly thermoregulation (Buchanan 2006). Anurans also have broad phenotypic, ecological, and behavioural diversity (Hödl and Amézquita 2001; Anderson and Wiens 2017; Moen 2019), which suggests that their visual systems may have adapted to contend with different light environments and functional demands. Several recent studies have investigated evolutionary correlations between species ecology and both morphological (eye size; Huang et al. 2019; Thomas et al. 2020; Shrimpton et al. 2021) and spectral (lens transmission and pigmentation; Yovanovich et al. 2020; Thomas et al. *in review*) features of anuran eyes. These studies found significant variation in anuran eye size and lens transmission that are associated with differences in behaviour and ecology suggesting substantial adaptation in visual function among anuran lineages. However, the molecular mechanisms underlying morphological and spectral adaptation in anuran visual systems have not yet been explored using a comparative evolutionary approach.

Here we focus on the molecular evolution of the visual opsin genes. These genes encode the protein component of visual pigments, the molecules contained in the photoreceptor cells of the retina that absorb light and initiate the phototransduction cascade that results in vision. In vertebrates there are ancestrally five visual opsin genes: one expressed in the dim-light sensitive, rod photoreceptors (*RH1*) and four expressed in spectrally distinct bright-light sensitive, cone photoreceptors (*LWS*, *RH2*, *SWS1*, *SWS2*). The different visual pigments formed by each of these opsins absorbs light maximally (λ_max_) at different wavelengths and these differences are controlled by the structure of the opsin protein as well as by the non-protein component of the visual pigment, the light-sensitive chromophore (Bowmaker 2008). Visual opsins have been independently lost and duplicated in many different lineages, resulting in as few as one visual opsin gene in some lineages, such as deep diving whales (Meredith et al. 2013), and up to 38 *RH1* copies in the spinyfin, *Diretmus argenteus* (Musilova et al. 2019). Further, variation in the sequences of opsin genes among species can result in considerable differences λ_max_ in among species. This variation in the number and type of visual opsins is one of the primary ways vertebrates can adapt their visual systems to different spectral environments (Loew and Lythgoe 1978; Bowmaker et al. 1994; Loew et al. 2002).

Shifts in spectral sensitivity of a particular visual opsin are termed spectral tuning and have been identified in all major vertebrate lineages (Yokoyama 2008; Davies et al. 2012). Spectral tuning can occur via changes to the opsin-coding sequence that result in the substitution of amino acid residues, particularly those lining the chromophore-binding pocket formed by the opsin’s seven transmembrane α-helices and alter the interaction between the opsin and the chromophore. Shifts in the spectral sensitivity of visual pigments can play an important role in the evolution, ecology, and behaviour of species. The most extreme example is in African lake cichlids where evidence suggests that divergent selection on spectral sensitivity in LWS drove speciation of two Lake Victoria cichlids through sensory drive (Seehausen et al. 2008). In neotropical cichlids, visual pigments have also been shown to be under divergent selective pressures associated with differences in habitat and light environments (Schott et al. 2014; Torres-Dowdall et al. 2015; Escobar-Camacho et al. 2017; Hauser et al. 2017; Hauser et al. 2021). In other vertebrates, similar associations between positive and divergent selection on opsin genes and shifts in light environments and behaviours have been found in diverse groups including in snakes (Schott et al. 2018), geckos (Schott et al. 2019), bats (Gutierrez et al. 2018a; Gutierrez et al. 2018b), whales (Dungan et al. 2016; McGowen et al. 2020), warblers (Bloch et al. 2015), and many other examples in teleost fishes (reviewed in Carleton et al. 2020).

In addition to spectral tuning, changes to the opsin sequence can also affect other aspects of visual pigment function including kinetic rates, such as light and thermal activation. For example, in the rod opsin (RH1) a D83N substitution has been identified as a potential dim-light adaptation by accelerating the formation of the active, signalling state of the visual pigment upon light activation (Sugawara et al. 2010). The effect of this mutation has been explored in a number of different groups that inhabit dim-light environments including cichlid fishes, bats, whales, echidnas, and bowerbirds (Sugawara et al. 2010; Bickelmann et al. 2012; van Hazel et al. 2016; Dungan and Chang 2017; Hauser et al. 2017). Like spectral tuning, these other functional properties of visual pigments may play an important role in visual adaptation but have been comparatively understudied.

Relative to other vertebrates, little is known about the diversity of photoreceptors and visual opsins in anurans and other amphibians. Four of the five visual opsin genes have been identified in anurans (*RH1*, *LWS*, *SWS1*, *SWS2*), but *RH2* has not been found in any amphibian and is presumed to have been lost early during their evolution (Bowmaker 2008; Schott et al. 2021a). These opsins may be found in as many as eight different photoreceptor types including two types of rods, one of which is unique to amphibians. The typical, RH1 rods (also called red rods) contain a green-absorbing pigment (λ_max_ of 491–503 nm; Table 1; Liebman and Entine 1968; Siddiqi et al. 2004) that is formed from rod opsin (RH1). The second, novel type of rod, historically (and confusingly) called a green rod, contains a blue-absorbing visual pigment (λ_max_ of 430-440 nm; Muntz and Reuter 1966; Dartnall 1967; Liebman and Entine 1968; Hisatomi et al. 1999; Darden et al. 2003; Govardovskii and Reuter 2014) that is formed from the SWS2 opsin typically expressed in cone photoreceptors. These SWS2 rods are rarer than the RH1 rods, but their proportion of the total rod population is highly variable in the limited number of species that have been studied to date (3–20%; Denton and Wyllie 1955; Nilsson 1964; Röhlich and Szél 2000), and this rod type may not be present in all frogs (e.g., *Oophaga pumilio;* Siddiqi et al. 2004). Further, the SWS2 opsin of at least some frogs, but none of the salamanders examined so far, have a unique amino acid residue, Thr47, that results in highly reduced thermal activation rates close to the level of RH1 opsins and much lower than any other cone opsins (Kojima et al. 2017).

**Table 1.** Maximum spectral sensitivity (λ_max_ in nm) of adult anuran photoreceptors estimated through microspectrophotometric (MSP) or electroretinographic (ERG) methodologies. Photoreceptors are grouped into rods and cones and then further divided based on λ_max_.

| Species | Rod 1 | Rod 2 | Cone 1 | Cone 2 | Cone 3 | Reference |
| --- | --- | --- | --- | --- | --- | --- |
| *Bufo bufo* | 502 | 432 |  |  |  | Govardovskii et al. 2000 |
| *Hyla cinerea* | 503 | 435 |  |  |  | King et al. 1993 |
| *Lithobates catesbeianus* | 502 | 432 | 570 |  | 433 | Hárosi 1982; Govardovskii et al. 2000 |
| *L. pipiens* | 502–503 | 432 | 575 | ~500 |  | Liebman and Entine 1968; Govardovskii et al. 2000 |
| *L. ridibunda* | 502 | 433 |  |  |  | Govardovskii et al. 2000 |
| *L. sphenoccephalus* | 501, 505 | ~437 | 579, 603 |  |  | Schott et al. 2021a |
| *Rana temporaria* | 501-503 | 434 | 562 |  | 431 | Koskelainen et al. 1994; Govardovskii et al. 2000 |
| *Oophaga pumilio* | 491 |  | 561 | 489 | 466 | Siddiqi et al. 2004 |
| *Rhinella marinus* | 503 | 432 |  |  |  | Govardovskii et al. 2000 |
| *Xenopus laevis* | 523–524 (A2) | 444–445 (A2) | 611 (A2) |  |  | Witkovsky et al. 1981; Govardovskii et al. 2000 |

Frogs also have at least three, and up to six, types of cones that include up to four different visual pigments. This includes red-sensitive LWS pigments (λ_max_ of ~560-575 nm; Liebman and Entine 1968; Liebman 1972), a green absorbing pigment spectroscopically indistinguishable from that in the RH1 rods (λ_max_ of ~500 nm), and a blue-absorbing pigment with a λ_max_ of ~430 nm (Liebman and Entine 1968; Hárosi 1982; Koskelainen et al. 1994). While the opsin identities of the visual pigments contained in all of the cones have not been determined, it seems highly likely that the green-sensitive cones contain the RH1 opsin also present in the RH1 rods, making this a rare example of the RH1 pigment being contained in a cone photoreceptor (Schott et al. 2016; de Busserolles et al. 2017). The blue cones could contain either SWS1 or SWS2 visual pigments and it is possible both types of cones are present, at least in some species. SWS1 expression has been detected in cones in both *Xenopus laevis* and in bullfrog (*Lithobates catesbeianus*; Hisatomi et al. 1998; Starace and Knox 1998), but direct evidence of SWS2 cones has not been found in frogs, unlike in salamanders (Isayama et al. 2014). Spectroscopically, three types of cones were identified in *Oophaga pumilio* (Siddiqi et al. 2004) that had λ_max_ of ~561, ~489, and ~466 nm. Only LWS is known to absorb maximally longer wavelengths (e.g., >550 nm), but the visual pigments in the 489 and 466 nm cones are less clear and could be some combination of RH1, SWS2, or SWS1.

To date, the photoreceptor and visual pigment complements of frogs have yet to be adequately resolved and almost no data are available on variation among species. Here we sequence visual opsins from 14 North American anuran species representing six families. We also take advantage of growing anuran genomic and transcriptomic resources to extract visual opsins from 14 species, which when combined with sequences available on Genbank, resulted in a total sample from 33 species and 12 families. Study species represent diverse evolutionary lineages and life histories and, thus we hypothesize they possess visual systems adapted to meet different ecological needs. We aim to: (1) determine which opsin genes are expressed in anuran retinas; (2) identify variation in opsin sequences among anuran species, including at potential spectral tuning and other functionally relevant sites; and (3) test for evidence of positive selection that may indicate functional adaptation to the distinct light environments inhabited by our study species.

## Methods

### Sample Collection

Thirteen of the fourteen anuran species newly sampled in this study are native to eastern Texas where they were collected. These include two species of “true toad” (*Incilius nebulifer* and *Anaxyrus woodhousii*); two species of chorus frog (*Pseudacris crucifer* and *P. fouquettei*); three species of treefrog (*Hyla chrysoscelis, H. versicolor*, and *H. cinerea*); four species of pond frog (*Lithobates catesbeianus, L. clamitans, L. palustris, and L. sphenocephalus*); one species of narrowmouth toad (*Gastrophryne carolinensis*); and one species of spadefoot toad (*Scaphiopus hurterii*). In addition to the thirteen native eastern Texas species, this study also includes the chirping frog *Eleutherodactylus cystignathoides*, which is introduced in eastern Texas, but native to the Rio Grande Valley in southern Texas. Our sampling also includes species for which genomic and transcriptomic resources are publicly available (see below). The phylogenetic relationships among study species are depicted in Figure 1.

**Figure 1.**
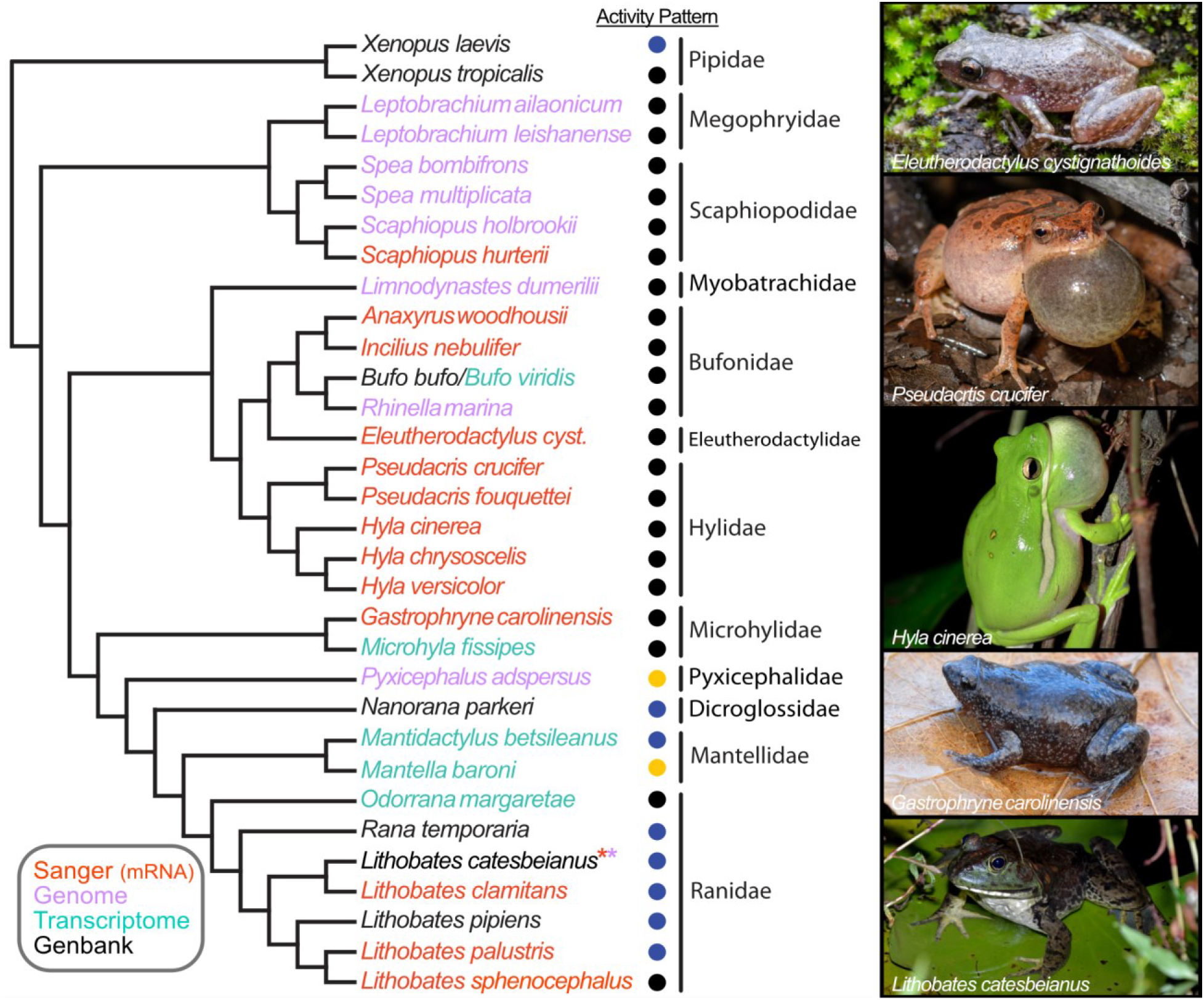
Phylogenetic tree illustrating evolutionary relationships among the study species based upon large-scale phylogenies (Pyron and Wiens 2011; Feng et al. 2017; Jetz and Pyron 2018; Streicher et al. 2018). The activity pattern of species is denoted with a circle where black = primarily nocturnal, yellow = primarily diurnal, and blue = both. The source of the sequence is also indicated through the colour of the species names. Sanger sequences were newly sequenced for the present study, while those from genomes and transcriptomes were newly extracted from existing assemblies. Sequences obtained from Genbank may have ultimately been derived from Sanger or whole genome sequencing. The individual species trees (with gene-specific sampling) and maximum likelihood gene trees used to analyze each gene can be found on Zenodo (Schott et al. 2021b). Photographs by MAK.

For the Texas frogs, up to five individuals per species were collected throughout the study period, from autumn of 2017 through spring of 2019. Most individuals were collected from ephemeral breeding ponds in the Stephen F. Austin Experimental Forest, which is part of the Angelina National Forest, and the adjacent Alazan Bayou Wildlife Management Area in southwestern Nacogdoches County, TX, USA. The strictly urban *E. cystignathoides* were collected on or near the Stephen F. Austin State University campus. All study animals were collected under permit and in compliance with the U.S. Forest Service, Texas Parks and Wildlife Department, and Nacogdoches city law enforcement. Following protocols described by the Herpetological Animal Care and Use Committee (2004) of the American Society of Ichthyologists and Herpetologists (ASIH) and approved by the SFASU Institutional Animal Care and Use Committee (Protocol # 2017-007), animals were euthanized via overdose of the anesthetic Tricaine methanesulfonate (MS-222). Euthanasia was confirmed prior to eye dissection by severing and pithing the spinal cord. Upon removal from the eye, each retina was immediately stored at −20°C in RNAlater (Thermo Fisher Scientific, Waltham, MA, USA).

### Opsin Sequencing

Total retinal mRNA was extracted from one of each study animal’s retinas with an RNeasy Mini Kit and QIAshredder (Qiagen, Valencia, CA, USA), quantified with a NanoVue spectrophotometer (GE Healthcare, Chicago, IL, USA), and stored at −80°C; the second retina remained in storage at −20°C. Standardized 0.4 μg mRNA aliquots were reverse transcribed using SuperScript™ IV Reverse Transcriptase (Invitrogen, Carlsbad, CA, USA) with an oligo(dT) primer to synthesize 20 μL aliquots of total cDNA. Fragments of each opsin-coding gene could then be amplified via polymerase chain reactions (PCR) for sequencing. Gene-specific and degenerate primers for anuran *RH1*, *LWS*, *SWS1*, and *SWS2* (Appendix 1) were designed using Primer3 (Rozen and Skaletsky 1999) from aligned GenBank reference sequences.

Each 25 μL PCR mixture contained 18.4 μL nuclease-free H_2_O, 2.0 μL 10X High Fidelity PCR Buffer, 1.0 μL 50 mM MgSO_4_, 0.5 μL mix of 10 mM-each dNTPs, 1.0 μL 10 μM forward primer, 1.0 μL 10 μM reverse primer, 0.1 μL Platinum *Taq* DNA Polymerase High Fidelity (5 U/μL) (Invitrogen, Carlsbad, CA, USA), and 1.0 μL template cDNA. Target gene fragments were amplified in a Mastercycler ep realplex thermocycler (Eppendorf, Hamburg, Germany) set to the following PCR profile: 95°C for 10 min; 94°C for 120 s; 35-50 cycles at 94°C for 30 s (denaturation), 45-50°C for 60 s (primer annealing), and 72°C for 120 s (polymerization); and 72°C for 120 s. Samples of PCR product were visualized with ethidium bromide in a 1% agarose gel to assess the effectiveness of each primer pair and to select suitable samples for cleanup and sequencing. PCR product was purified with the Wizard^®^ SV Gel and PCR Clean-Up System (Promega Corporation, Madison, WI, USA), quantified, and prepared according to specifications set by the DNA Sequencing Facility at the University of Texas at Austin for nucleotide sequencing via the chain-termination method (Sanger et al. 1977). Returned partial sequences were identified to the gene via nucleotide BLAST (Altschul et al. 1990). Prior to further analysis, partial sequences of the same gene from the same species were cleaned and merged into a consensus sequence in Geneious 10 (Biomatters, Ltd., Auckland, New Zealand; Kearse et al. 2012). In the case of *Lithobates clamitans*, only one of the two individuals collected was sequenced. Among other species, opsins were sequenced from two individuals in *Incilius nebulifer*, *Eleutherodactylus cystignathoides*, *Hyla chrysoscelis*, *H. versicolor*, *Gastrophryne carolinensis*, and *L. clamitans;* three individuals in *Anaxyrus woodhousii*, *H. cinerea*, *Pseudacris fouquettei*, *L. sphenocephalus*, and *L. palustris*; and four individuals in *P. crucifer* and *Scaphiopus hurterii*.

### Visual Opsin Gene Datasets

Additional visual opsin sequences were obtained from the NCBI Genbank database and were extracted from all available anuran genome and transcriptome assemblies using BLAST (Additional File 1). We also assembled *Mantidactylus betsileanus* transcriptome reads (from Wollenberg Valero et al. 2017) *de novo* using Trinity (Grabherr et al. 2011) and extracted visual opsin coding regions from the resulting assembly. Total numbers of sequences obtained for each opsin, and their source, are outlined in Additional File 1.

Coding regions for each of the four visual opsin genes obtained from anurans (*RH1*, *LWS*, *SWS1*, *SWS2*) were aligned using MUSCLE codon alignment as implemented in MEGA (Edgar 2004; Tamura et al. 2011) followed by manual correction. Maximum likelihood (ML) gene trees were inferred for each gene using PhyML 3 (Guindon et al. 2010) under the GTR+G+I model with a BioNJ starting tree, the best of NNI and SPR tree improvement, and aLRT SH-like branch support (Anisimova and Gascuel 2006). In additional to the ML topologies, trees were modified to match the expected species relationships based on the large-scale phylogenies of Pyron and Wiens (2011), Feng et al. (2017), Jetz and Pyron (2018), and Streicher et al. (2018) to produce species tree topologies.

### Selection Analyses

To estimate the strength and form of selection acting on the visual opsin genes in anurans, each dataset was analyzed using codon-based likelihood models from the codeml program of the PAML 4 software package (Yang 2007). Specifically, we used the random sites models (M0, M1a, M2a, M2a_rel, M3, M7, M8a, and M8) to infer alignment-wide selection patterns and to test for positive selection acting on any of the genes. All analyses were run with varying starting values to avoid potential local optima. To determine significance, model pairs were compared using a likelihood ratio test (LRT) with a χ^2^ distribution. Analyses were run using both the ML gene tree and species tree topologies modified to contain the basal trichotomy required by PAML. The Bayes Empirical Bayes (BEB) approach was used to identify individual sites with a high posterior probability of being in the positively selected class of sites.

We also analyzed the data using the HYPHY model FUBAR (Pond and Frost 2005; Murrell et al. 2013) implemented on the Datamonkey webserver (Delport et al. 2010). This model uses a hierarchical Bayesian method to average over a much larger number of site classes than the PAML models and importantly allows for an independently estimated value for dS.

## Results

### Frog Visual Opsins

Partial coding sequences of four opsins—RH1, LWS, SWS1, and SWS2—were recovered from the retinal mRNA of fourteen anuran species (Table 2, Additional File 1). Several primer pairs were unsuccessful, and we failed to amplify sequences, or parts of sequences, from a number of species (Table 2, Additional File 1). Consequently, we do not consider the lack of recovery of any of the opsins genes from retinal mRNA as evidence for a lack of expression or gene loss. Additional coding sequences were extracted from available frog genomes and transcriptomes, as well from Genbank. This resulted in 31 RH1, 28 LWS, 26 SWS1, and 30 SWS2 sequences in total (Table 2, Additional File 1).

**Table 2.**
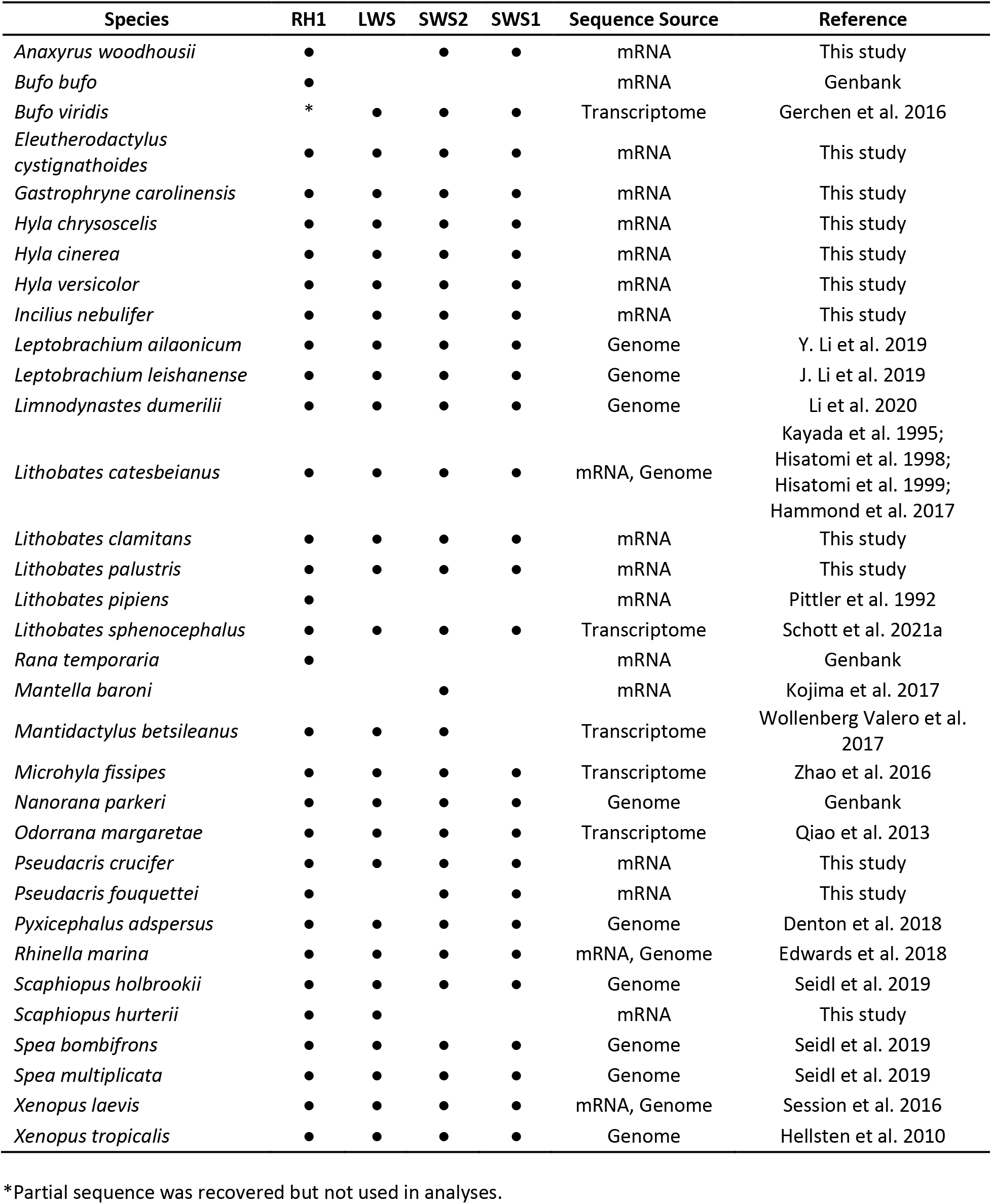
Summary of visual opsin genes sequenced or extracted in the current study. Full details, including individual accession numbers can be found on Zenodo (Schott et al. 2021b).

| Species | RH1 | LWS | SWS2 | SWS1 | Sequence Source | Reference |
| --- | --- | --- | --- | --- | --- | --- |
| *Anaxyrus woodhousii* | ● |  | ● | ● | mRNA | This study |
| *Bufo bufo* | ● |  |  |  | mRNA | Genbank |
| *Bufo viridis* | * | ● | ● | ● | Transcriptome | Gerchen et al. 2016 |
| *Eleutherodactylus cystignathoides* | ● | ● | ● | ● | mRNA | This study |
| *Gastrophryne carolinensis* | ● | ● | ● | ● | mRNA | This study |
| *Hyla chrysoscelis* | ● | ● | ● | ● | mRNA | This study |
| *Hyla cinerea* | ● | ● | ● | ● | mRNA | This study |
| *Hyla versicolor* | ● | ● | ● | ● | mRNA | This study |
| *Incilius nebulifer* | ● | ● | ● | ● | mRNA | This study |
| *Leptobrachium ailaonicum* | ● | ● | ● | ● | Genome | Y. Li et al. 2019 |
| *Leptobrachium leishanense* | ● | ● | ● | ● | Genome | J. Li et al. 2019 |
| *Limnodynastes dumerilii* | ● | ● | ● | ● | Genome | Li et al. 2020 |
| *Lithobates catesbeianus* | ● | ● | ● | ● | mRNA, Genome | Kayada et al. 1995;<br>Hisatomi et al. 1998;<br>Hisatomi et al. 1999;<br>Hammond et al. 2017 |
| *Lithobates clamitans* | ● | ● | ● | ● | mRNA | This study |
| *Lithobates palustris* | ● | ● | ● | ● | mRNA | This study |
| *Lithobates pipiens* | ● |  |  |  | mRNA | Pittler et al. 1992 |
| *Lithobates sphenoccephalus* | ● | ● | ● | ● | Transcriptome | Schott et al. 2021a |
| *Rana temporaria* | ● |  |  |  | mRNA | Genbank |
| *Mantella baroni* |  |  | ● |  | mRNA | Kojima et al. 2017 |
| *Mantidactylus betsileanus* | ● | ● | ● |  | Transcriptome | Wollenberg Valero et al. 2017 |
| *Microhyla fissipes* | ● | ● | ● | ● | Transcriptome | Zhao et al. 2016 |
| *Nanorana parkeri* | ● | ● | ● | ● | Genome | Genbank |
| *Odorrana margaretae* | ● | ● | ● | ● | Transcriptome | Qiao et al. 2013 |
| *Pseudacris crucifer* | ● | ● | ● | ● | mRNA | This study |
| *Pseudacris fouquettei* | ● |  | ● | ● | mRNA | This study |
| *Pyxicephalus adspersus* | ● | ● | ● | ● | Genome | Denton et al. 2018 |
| *Rhinella marina* | ● | ● | ● | ● | mRNA, Genome | Edwards et al. 2018 |
| *Scaphiopus holbrookii* | ● | ● | ● | ● | Genome | Seidl et al. 2019 |
| *Scaphiopus hurterii* | ● | ● |  |  | mRNA | This study |
| *Spea bombifrons* | ● | ● | ● | ● | Genome | Seidl et al. 2019 |
| *Spea multiplicata* | ● | ● | ● | ● | Genome | Seidl et al. 2019 |
| *Xenopus laevis* | ● | ● | ● | ● | mRNA, Genome | Session et al. 2016 |
| *Xenopus tropicalis* | ● | ● | ● | ● | Genome | Hellsten et al. 2010 |

\*Partial sequence was recovered but not used in analyses.

For each of the available frog genomes all four expected visual opsins (*RH1*, *LWS*, *SWS1*, and *SWS2*) were recovered. In the *Pyxicephalus adspersus* genome we identified two *LWS* genes, one on each of the two sex chromsomes (W and Z). The two sequences are relatively divergent sharing 93.5% amino acid identity (91.8% nucleotide identity) and the Z chromosome sequence has a single amino acid deletion at site 331 (note site numbering is relative to bovine rhodopsin throughout). Phylogenetic analyses revealed the sequences are most closely related to each other suggesting that they are a species-specific (or at least lineage-specific) duplication (Figure 2).

**Figure 2.**
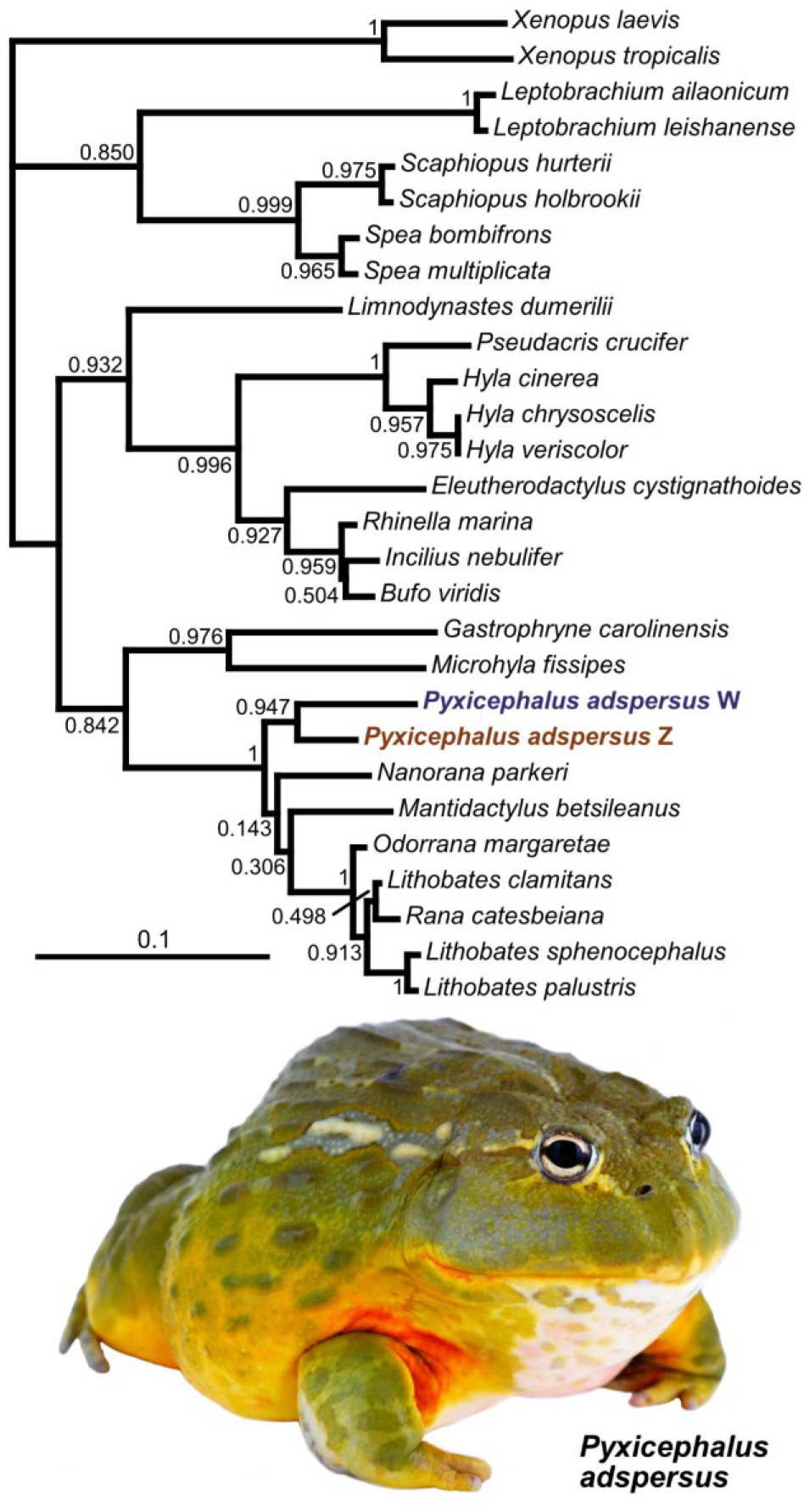
Maximum likelihood gene tree for *LWS* depicting the two LWS genes in *Pyxicephalus adspersus*. The gene tree was inferred using PhyML 3 (Guindon et al. 2010) under the GTR+G+I model with a BioNJ starting tree, the best of NNI and SPR tree improvement. Branch support values (aLRT SH-like; Anisimova and Gascuel 2006) are shown at the nodes. The basal trichotomy is required by PAML and was manually created. Photograph by John Clare.

### Variation at Known Spectral Tuning Sites

Each of the four visual opsins possessed at least one amino acid substitution at a gene-specific site known in other vertebrates to tune spectral sensitivity of visual pigments (Table 3). The RH1 gene exhibited a change from the nonpolar, aliphatic amino acid alanine (A) to the polar, uncharged serine (S) at position 299 (notated as A299S; Table 3, Schott et al. 2021b) in several species (*Bufo bufo*, *Microhyla fissipes*, *Pseduacris crucifer*, *Rhinella marina*, *Scaphiopus hurterii*, *Sc. holbrookii*, *Spea bombifrons*, *and Sp. multiplicata*). This change is responsible for a slight (2 nm) shift in bovine and cetacean RH1 (Dungan and Chang 2017) and has been implicated in spectral tuning in deep dwelling teleost fishes (Hunt et al. 1996; Hunt et al. 2001). The substitution Y102F was found in *Leptobrachium ailaonicum* and *Le. leishanense*. This change may produce a slight blue-shift, perhaps in combination with another change not found in frogs (Y96V; Yokoyama 2008). The substitution L194P occurs in *Microhyla fissipes*. This site has been identified as a spectral tuning site in RH1, but the documented substitution is P194R and it may only have an effect in combination with other residues (Yokoyama 2008). Additionally, anuran RH1 varied at six amino acid positions (46, 52, 93, 97, 109, 116) known to affect the spectral sensitivity of other vertebrate visual pigments (Table 3).

**Table 3.** Variation in anuran opsin sequences at known spectral tuning sites (based on those identified in Yokoyama 2008). The residues we identified in anurans are listed for each spectral tuning site while those sites with variation in the same opsin are bolded.

| Site (RH1 Numbering) | Known From | Known Spectral Variants | Residues in Anurans |  |  |  |
| --- | --- | --- | --- | --- | --- | --- |
|  |  |  | RH1 | SWS1 | SWS2 | LWS |
| 44 | SWS2 | M/T | M | M | M | M |
| 46 | SWS1/2 | F/T/L | L/M | **V/M/A/F** | F | F |
| 49 | RH2, SWS1 | S/F/A/V/L | L | **L/I/F/V** | I | A/I/G/L/F/S |
| 52 | RH2, SWS1 | L/M/T/F | F/L | **T/A** | F | V/C/I |
| 83 | RH1/2 | D/N | N | G | N | D |
| 86 | RH2, SWS1 | M/T/F/S/L/Y | M | **M/I** | V | E |
| 90 | SWS1 | S/C | G | S | G | A |
| 91 | SWS1/2 | V/I/S/P | F | **I/N** | S | S |
| 93 | SWS1 | T/P/L/I | I/V | **T/I/V/P** | T/V/M | I |
| 94 | SWS2 | A/S/C | T | V | A | S |
| 96 | RH1 | Y/V | Y | V/I/M | Y | F/I/A/V/C |
| 97 | RH2, SWS2 | T/A/S/C | T/S | S/N | S | N |
| 102 | RH1 | Y/F | **Y/F** | Y/C | Y | Y |
| 109 | SWS1/2 | V/A/G | G/T | **V/A/F/T** | A | L/M |
| 113 | SWS1 | E/D | E | **E** | E | E |
| 114 | SWS1 | A/G | G | **G/A** | G | G |
| 116 | SWS1/2 | L/V/T | F/C | **V/I/T** | T | T |
| 118 | SWS1/2 | S/T/A/G | T | **T/S** | T | S/A |
| 122 | RH1, SWS1/2 | E/I/Q/M | E | **L** | **M/I** | I |
| 124 | RH1 | A/S/G/V | A | T/I | S/G | G/A |
| 132 | RH1 | A/S | A | A | A | A |
| 164 | RH2, LWS | S/A | A | G | G/S/A | **A/S** |
| 181 | LWS | H/Y | E | E | E | H |
| 194 | RH1 | P/R | **L/P** | V/I | V | G |
| 195 | RH1 | N/A | K | G | N | S/N |
| 207 | RH2 | M/L | M | I/V | M/I/L | L |
| 208 | RH1 | F/Y | F | F | F | M |
| 211 | RH1 | H/C | H | C | C | C |
| 261 | RH1, SWS2, LWS | F/Y | F | F | F | Y |
| 265 | SWS2 | W/Y | W | Y | W | W |
| 269 | SWS2, LWS | A/S/T | A | A | A | T |
| 292 | RH1/2, SWS2, LWS | A/S | A | A | S | A |
| 295 | RH1 | A/S | A | S | S | A |
| 299 | RH1 | A/S | **A/S** | C | T | T |
| 300 | RH1 | I/T/L | I | V | V | I |

On the LWS opsin, an amino acid change occurred at known LWS tuning site 164 (anuran LWS-specific site 179), at which position *B. viridis*, *E. cystignathoides*, *I. nebulifer*, *Ma. betsileanus*, *N. parkeri*, *Odorrana margaretae*, *Pyxicephalus adspersus*, *R. marina*, *Sc. hurterii*, *Sc. holbrookii*, *Sp. bombifrons*, *Sp. multiplicata*, and *X. tropicalis* expressed alanine, and remaining species expressed serine (Table 3, Schott et al. 2021b). The substitutions A164S and S164A were shown to shift λ_max_ by 6 and −7 nm, respectively, in mammalian LWS (Asenjo et al 1994; Yokoyama et al. 2005, 2008). Anuran LWS also varied at three RH1 tuning sites (96, 124, and 195), two RH2/SWS1 tuning sites (49 and 52), and two SWS1/2 tuning sites (109 and 118), many of which include non-conservative amino acid substitutions and known spectral variants that could be expected to effect λ_max_ (Table 3).

SWS1 exhibited the greatest number of amino acid changes at gene-specific tuning sites (Table 3, Schott et al. 2021b). All 10 variable SWS1-specific sites (46, 49, 52, 86, 91, 93, 109, 114, 116, and 118) occurred within the first three transmembranes. At site 46 (anuran SWS1-specific site 42), the species expressed one of four residues, although none of these includes the known SWS1 spectral variant (F46T; Table 3; Yokoyama 2008). Site 49, which varied among four residues in our sample (L, I, F, V), did include the known spectral variants F49V/L (Table 3).

The substitutions F49V and F49L are responsible for a shift from UV λ_max_ (~360 nm) to violet λ_max_ (309+ nm) in combination with substitutions at several other sites (Yokoyama 2008). Sites 52, 86, and 91 were less variable and did not have known variants (Table 3). There were four residues found at site 93 (T, I, V, P) that include known spectral variants T93P and I93T (Table 3). Only four species had P93 (*X. laevis*, *X. tropicalis*, *Le. leishanense*, and *M. fissipes*, three had I (*Lithobates palustris*, *Li. sphenocephalus* and *N. parkeri*), five have V, and the rest T (Table 3). The T93P substitution was shown to contribute to the red-shifted λ_max_ of *X. laevis* SWS1 but may have little effect in isolation (Takahashi and Yokoyama 2005). The substitution I93T was shown to cause a −6 nm shift (Yokoyama et al. 2005). The effects of the other residues found in anurans at this site are not known. Site 109 had four variants in anurans (V, A, F, T). The substitution V109A was also identified as contributing to the violet λ_max_ of *X. laevis* SWS1, but similarly in isolation had no effect (Yokoyama et al. 2005). At site 114 two variants were found (A and G) and the substitution A114G was shown to result in a 5 nm shift in an inferred ancestral SWS1 pigment (Shi and Yokoyama 2003). Sites 116 and 118 varied at three (V, I, T) and two (T, S) sites, respectively, and substitutions at both sites contribute to the red-shifted λ_max_ of *X. laevis* SWS1 in coordination with substitutions at other sites, but were not found to have individual effects (Takahashi and Yokoyama 2005). Finally, in addition to variation at the aforementioned tuning sites, anuran SWS1 also varied at known RH1 tuning sites 96, 102, 124, 194, RH2 site 207, and RH2/SWS2 tuning site 97 (Table 3).

On the SWS2 opsin, amino acid variation occurred at gene-specific tuning site 122 (anuran SWS2-specific site 131), with *G. carolinensis*, *Le. ailaonicum* and *Le. leishanense*, *M. fissipes*, *Sc. hurterii*, *Sc. holbrookii*, *Sp. bombifrons*, *Sp. multiplicata*, *X. laevis*, and *X. tropicalis* expressing isoleucine and remaining species expressing methionine (Table 3, Schott et al. 2021b). The substitution I122M resulted in a −6 nm shift in newt (*Cynops pyrrhogaster*) SWS2 (Takahashi and Ebrey 2003). In addition, anuran SWS2 varied at four amino acid positions (93, 124, 164, and 207) known in other opsins to affect spectral sensitivity (Table 3).

### Variation at other functionally relevant sites

RH1 site 83 has been suggested to be associated with dim-light adaptation through a D83N substitution (Sugawara et al. 2010). The anurans we sampled all had N83. S299A (and vice versa) was found to affect retinal release rate in mammals (Dungan et al. 2017) and the sampled anurans varied among these two residues. Additionally, all frogs in our sample have SWS2 with T47, a mutation that was shown to result in increased dark state stability (low thermal isomerization rate; Kojima et al. 2017). Other sites known to affect kinetic rates, such as RH1 sites 59, 288, and 292 (Castiglione et al. 2017; Dungan and Chang 2017) were conserved in our sample of frogs.

In all four opsins, amino acid changes also occurred at additional sites forming the chromophore-binding pocket (list of sites provided in (Hunt et al. 2001). These included two sites (54 and 119) on RH1, two sites (119 and 160) on LWS, six sites (47, 82, 120, 258, 271, and 307) on SWS1, and two sites (207 and 258) on SWS2. Variation at site 119 included polarity changes in both RH1, with the amino acid change L119V in *G. carolinensis* and *M. fissipes*, and LWS, with the change V119T in several species (Table 3, Schott et al. 2021b). Another polarity change occurred at LWS site 160, at which few species have a S160A substitution. Of the six variable sites lining the chromophore-binding pocket in SWS1, only one included a polarity change where species varied between S, T, and A at site 120.

### Site-Specific Positive Selection

Using the PAML M8 model we found statistically significant positive selection at a small proportion of sites in both anuran RH1 and LWS with both the ML gene tree and species tree topologies (Tables 4 and 5, Schott et al. 2021b). Four RH1 sites were inferred to be under positive selection with a BEB posterior probability of >80% (39, 107, 213, 270; Table 6). None of those sites have previously been identified to affect spectral tuning, but most are near known sites. FUBAR analysis identified one of the same sites as M8 (213) in addition to five other sites (65, 97, 112, 169, 277; Table 6, Schott et al. 2021b). BEB analyses of the PAML M8 model inferred two LWS sites to be under positive selection with posterior probability >80% (49, 217), while FUBAR identified three (49, 154, 166; Table 6). One of these (49) is a known spectral tuning site in the RH2 and SWS1 opsins. No evidence of positive selection in SWS1 or SWS2 was detected with the PAML models, although two sites were identified with greater than 90% posterior probability in SWS1 and one site in SWS2 using FUBAR (Table 6, Schott et al. 2021b).

**Table 4.**
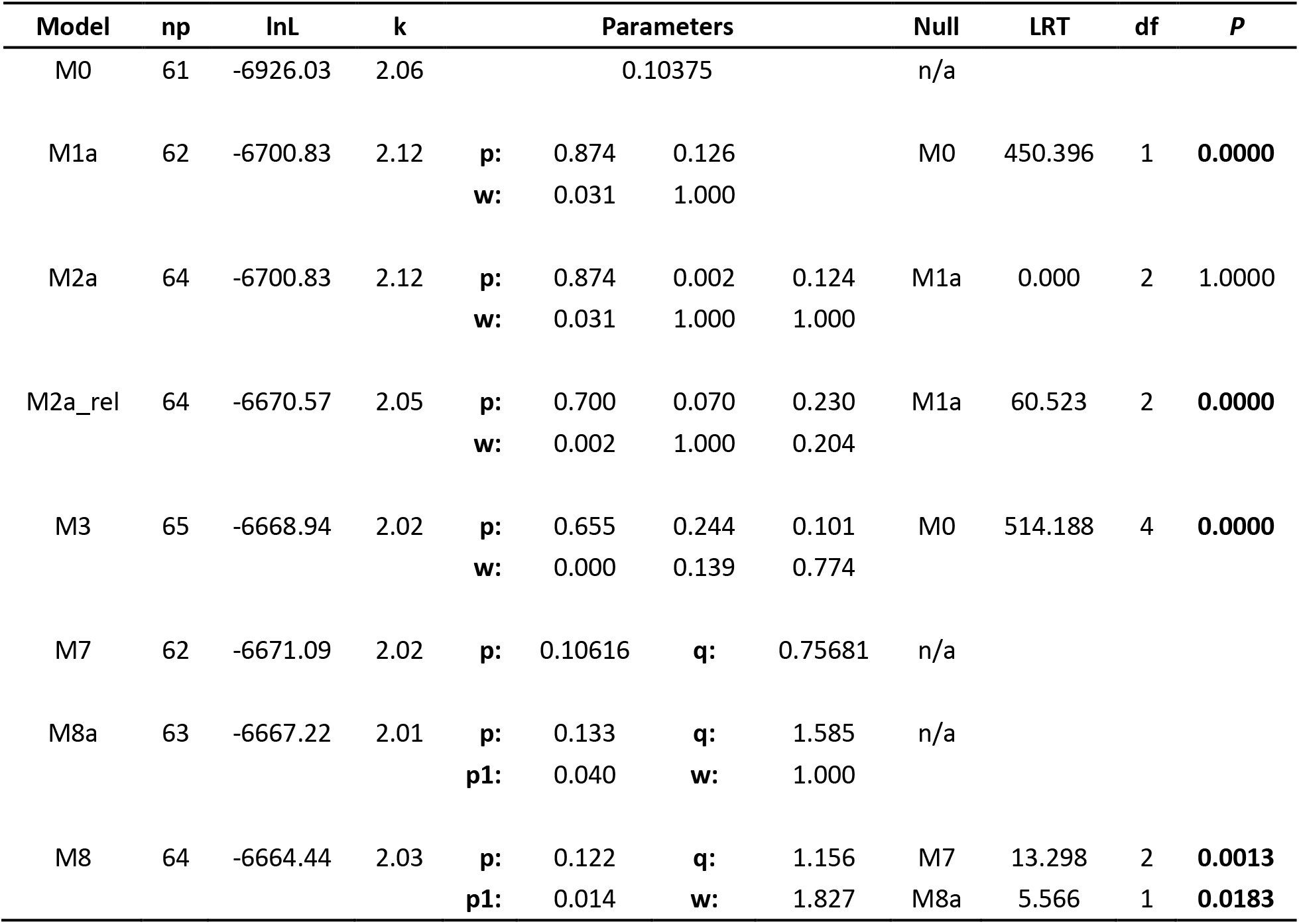
Results of PAML analyses performed on RH1 using the species topology. Results using the RH1 ML gene tree are similar and can be found on Zenodo (Schott et al. 2021b).

**Table 5.** Results of PAML analyses performed on LWS using the species topology. Results using the LWS ML gene tree are similar and can be found on Zenodo (Schott et al. 2021b).

| Model | np | lnL | k | Parameters |  |  |  | Null | LRT | df | P |
| --- | --- | --- | --- | --- | --- | --- | --- | --- | --- | --- | --- |
| M0 | 55 | -7462.48 | 2.13 | 0.09662 |  |  |  | n/a |  |  |  |
| M1a | 56 | -7280.33 | 2.23 | **p:** | 0.867 | 0.133 |  | M0 | 364.292 | 1 | **0.0000** |
|  |  |  |  | **w:** | 0.033 | 1.000 |  |  |  |  |  |
| M2a | 58 | -7278.72 | 2.24 | **p:** | 0.866 | 0.131 | 0.003 | M1a | 3.227 | 2 | 0.1992 |
|  |  |  |  | **w:** | 0.034 | 1.000 | 3.590 |  |  |  |  |
| M2a_rel | 58 | -7228.79 | 2.08 | **p:** | 0.203 | 0.031 | 0.766 | M1a | 103.075 | 2 | **0.0000** |
|  |  |  |  | **w:** | 0.320 | 1.000 | 0.012 |  |  |  |  |
| M3 | 59 | -7228.61 | 2.08 | **p:** | 0.770 | 0.206 | 0.024 | M0 | 467.743 | 4 | **0.0000** |
|  |  |  |  | **w:** | 0.013 | 0.338 | 1.159 |  |  |  |  |
| M7 | 56 | -7232.66 | 2.08 | **p:** | 0.14811 | **q:** | 1.13691 | n/a |  |  |  |
| M8a | 57 | -7229.25 | 2.08 | **p:** | 0.174 | **q:** | 1.866 | n/a |  |  |  |
|  |  |  |  | **p1:** | 0.024 | **w:** | 1.000 |  |  |  |  |
| M8 | 58 | -7227.31 | 2.09 | **p:** | 0.159 | **q:** | 1.373 | M7 | 10.712 | 2 | **0.0047** |
|  |  |  |  | **p1:** | 0.004 | **w:** | 2.476 | M8a | 3.878 | 1 | **0.0489** |

**Table 6.** Opsin amino acid sites inferred to be under positive selection with at least 80% posterior probability by either the BEB analyses of M8 model or with FUBAR. Sites numbers are relative to those in bovine RH1. Full PAML and FUBAR results tables can be found on Zenodo (Schott et al. 2021b).

| Opsin | Site Number | PAML M8 BEB |  | FUBAR |  |
| --- | --- | --- | --- | --- | --- |
| | | Posterior Probability | $\omega$ | Posterior Probability | $\omega$ |
| RH1 | 39 | 0.988 | $1.498 \pm 0.106$ | 0.791 | 3.272 |
| RH1 | 65 | - | - | 0.865 | 3.005 |
| RH1 | 97 | - | - | 0.862 | 2.777 |
| RH1 | 107 | 0.979 | $1.192 \pm 0.124$ | 0.019 | 0.399 |
| RH1 | 112 | - | - | 0.92 | 4.158 |
| RH1 | 169 | 0.753 | $1.314 \pm 0.339$ | 0.987 | 8.027 |
| RH1 | 213 | 0.964 | $1.482 \pm 0.145$ | 0.94 | 5.555 |
| RH1 | 270 | 0.842 | $1.387 \pm 0.283$ | 0.15 | 0.878 |
| RH1 | 277 | - | - | 0.885 | 3.475 |
| LWS | 49 | 0.995 | $1.524 \pm 0.238$ | 0.917 | 6.790 |
| LWS | 154 | 0.551 | $1.153 \pm 0.401$ | 0.918 | 4.358 |
| LWS | 166 | - | - | 0.864 | 2.830 |
| LWS | 217 | 0.898 | $1.442 \pm 0.254$ | 0.543 | 2.240 |
| LWS | 49 | - | - | 0.917 | 6.790 |
| LWS | 154 | - | - | 0.918 | 4.358 |
| LWS | 166 | 0.864 | 2.830258 | 0.864 | 2.830 |
| SWS1 | 120 | - | - | 0.909 | 7.280 |
| SWS1 | 159 | - | - | 0.906 | 3.560 |
| SWS2 | -2 | - | - | 0.901 | 7.194 |

## Discussion

Using a combination of retinal cDNA sequencing and previously published genomic and transcriptome resources we obtained visual opsin genes for 33 anuran species spanning 12 families. We found that anurans generally possess four of the visual opsins common to vertebrates (*RH1*, *LWS*, *SWS1*, *SWS2*) with no evidence of the *RH2* opsin gene. While we had variable recovery of opsins from retinal cDNA, we did not find any evidence for loss of visual opsins in any of the species for which genomic data were available. We identified a single gene duplication, in *Pyxicephalus adspersus*, where a distinct *LWS* gene was found on each of the two sex chromosomes (Z and W). Overall we found considerable variation in each of the four opsins across anurans at both previously known and potentially new functional sites. In addition, we found evidence for positive selection in all four genes at a small subset of sites, most notably in *RH1* and *LWS*. Below we discuss these findings in terms of how they may affect spectral tuning and dim-light adaptation in anurans that inhabit different light environments.

### Spectral Tuning Variation in Anuran Visual Opsins

We identified a large amount of variation in each of the four visual opsins at known spectral tuning sites. However, much of this variation was between residues not found, or at least not explored, in other groups making it difficult to predict the effect of the differences in protein sequence. This is further complicated by the relative lack of data on visual pigment spectral absorbances available for anurans (Table 1), which limits our ability to infer the effect of particular substitutions on the spectral absorbance of the visual pigment. Additionally, in each opsin, considerable variation was found at spectral tuning sites that are known from other visual opsins. While some of these sites are likely to affect spectral tuning in multiple visual opsins, others will have a more restricted effect due to interactions with other residues in the protein. Despite this, our results highlight that there is likely considerable unappreciated variation in the spectral absorbances of anuran visual pigments and we have identified numerous candidates for further functional studies.

Based on the limited available data, the RH1 visual pigment of most frogs, including *Lithobates* spp., *Bufo* spp., and *Hyla cinerea*, are reported to have a λ_max_ of ~502 nm. Exceptions to this are *Oophaga pumilio* with a λ_max_ of 491 nm and *X. laevis* with a λ_max_ of 535 nm (Table 1). Unfortunately, we did not have an *O. pumilio* sample (or other dendrobatid) to evaluate potential causes of the blue-shifted λ_max_. In *X. laevis*, the red-shifted λ_max_ is caused by the use of a different chromophore that is derived from vitamin A2 (3,4-didehydroretinal, referred to as A2), as opposed to the more typical A1 chromophore (retinal) used by most vertebrates. The A2 chromophore is found in some anuran tadpoles but is replaced by A1 chromophore in the adults of most frog species, whereas other frogs exclusively use A1 (Bridges 1972). In *X. laevis*, however, A2 is used throughout its lifecycle (Bridges et al. 1977). Partial replacement of A2 by A1 and an A1 variant (9-*cis* retinal) resulted in λ_max_ of *X. laevis* RH1 visual pigment of 511 and 498 nm (Witkovsky et al. 1981) suggesting that the RH1 opsin may be slightly blue-shifted relative to most other known frog RH1s. *Xenopus laevis* RH1 did not differ in any known RH1 tuning sites from the other species in our dataset with measured λ_max_ (e.g., *Lithobates* spp.), but does differ at a tuning site known from SWS1 (site 93) making this a potential spectral tuning site candidate. Additionally, they differ at 5 of the 9 sites identified as being positively selected in RH1 (sites 39, 107, 112, 169, 213) also suggesting these sites may influence spectral tuning, while the other positively selected sites may influence other aspects of visual pigment function. In particular, Q107P and L213T may be of particular interest for future studies.

The LWS cones of anurans, again based on limited data, have variable spectral sensitivities ranging from λ_max_ of 561-579 nm for A1-based pigments (Table 1). Unfortunately, sequences for *O. pumilio* and *R. temporaria*, which are reported to have blue-shifted λ_max_ around ~561 nm are not available, but the LWS-specific spectral tuning substitution S164A likely contributes to this shift. Species with λ_max_ ≥570 nm (e.g., *Lithobates catesbaenia* and *L. sphenocephalus*) have S164 and the substitution S164A was shown to shift λ_max_ −7 nm when mutated in human LWS (Asenjo et al. 1994). However this substitution alone is not enough to account for the difference and thus substitutions at other sites are likely to also affect LWS spectral tuning in anurans. The four sites identified in anuran LWS as being positively selected are likely also to play a role, especially site 49, which was highly variable and is known to effect spectral tuning in other visual opsins.

Evidence for SWS1 cones in anurans was previously very limited (Hisatomi et al. 1998; Starace and Knox 1998). While our data cannot inform on potential combinations of visual pigments in different types of cones, that fact that *SWS1* does not appear to have been lost in any species, and is under high selective constraint, suggests that SWS1 visual pigment is present in anuran photoreceptors, at least at some point in their life cycle. This further suggests that SWS1 cones are common among anurans and are just difficult to detect with methods such as microspectrophotometry (MSP) and electroretinograms (ERG). A potential convergence of the λ_max_ of SWS1 and SWS2 (see above) may further complicate this, although in *X. laevis* the λ_max_ of these pigments expressed *in vitro* differed by 9 nm (425 vs 434 nm, respectively; Starace and Knox 1998; Darden et al. 2003). It is also possible that SWS1 is co-expressed with another opsin as is the case in the cones of salamanders and several other vertebrates (Dalton et al. 2014; Isayama et al. 2014), but an opsin that is only co-expressed would be novel. Another possibility is that, in some anuran species, SWS1 is only expressed at certain life stages, such as in tadpoles. Ontogenetic shifts in expression of visual opsins is fairly common in teleost fishes (Carleton et al. 2020), but the only study of expression profiles in a frog (*L. sphenocephalus*) found that SWS1 was expressed at a low, but consistent level in both tadpoles and post-metamorphic juvenile frogs (Schott et al. 2021a).

Overall, the current literature suggests anuran SWS1 λ_max_ is fairly conserved and varies only between 425 and 433 nm, and yet our molecular data showed that SWS1 was the most variable of the four visual opsins at known spectral tuning sites. While this high sequence diversity perhaps indicates more variation in λ_max_ than is currently documented, we found that anuran SWS1 was under high selective constraint and had little evidence of positively selected sites. Thus, potential spectral shifts may have only occurred a small number of times, in specific lineages, which would not leave a signature of positive diversifying selection detectable by the codon models we employed. Estimating the effect on spectral tuning of variation at known spectral tuning sites remains challenging because many of the sites have interacting effects and, in some cases, the specific residues found in anurans are not found in other groups (Takahashi and Yokoyama 2005; Hauser et al. 2014). Finally, studies of SWS1 λ_max_ in anurans to date have not yet found evidence that spectral sensitivity of this visual opsin is shifted into the UV. Shifts between violet and UV sensitivity are relatively common in vertebrates, especially in birds where data supports at least 14 shifts between violet and UV sensitivity (Ödeen and Håstad 2013), although the evolutionary drivers of these shifts are not well understood. While several of the changes we identified suggest UV-sensitivity of SWS1 in anurans may be possible, further functional studies will be required to answer this question.

Uniquely in anurans and some salamanders the SWS2 opsin is expressed in SWS2 rods (also known as ‘green’ rods). In salamanders, the SWS2 opsin is also expressed in cones but direct evidence of SWS2 cones is lacking in anurans (Isayama et al. 2014; Hisatomi et al. 1993, Darden et al. 2003; but see Siddiqi et al. 2004). The λ_max_ of anuran SWS2 rods, at least based on current data, is conserved around ~432 nm (Table 1). *Lithobates catesbaeianus* and *R. tempororia* also have cones with the same λ_max_ as the SWS2 rods, although immunohistochemical evidence in *L. catesbaeianus* shows no evidence of SWS2 expression in cones, suggesting that the SWS1 and SWS2 pigments may have converged on the same λ_max_ (Hárosi 1982; Koskelainen et al. 1994; Donner and Yovanovich 2020). The λ_max_ of SWS2 rods in *X. laevis* was estimated to be 445 nm with A2 (Witkovsky et al. 1981), but when the SWS2 pigment was expressed *in vitro* with A1 chromophore the λ_max_ was found to be 434 nm closely matching the absorbances in other species estimated from intact SWS2 rod photoreceptors. *Xenopus laevis* and the other species for which SWS2 λ_max_ has been estimated (e.g., *L. catesbaeianus, Bufo bufo*; Table 1) differ at the SWS2 spectral tuning site 122 (I in *X. laevis*, M in the others). In the newt *Cynops pyrrhogaster* I122M resulted in a −6 nm shift (Takahashi and Ebrey 2003), which suggests that the spectral tuning effect of this site may differ between anurans and salamanders, although the possibility of compensatory mutations in anurans cannot be ruled out. *Xenopus laevis* and the other species also differed at a number of spectral tuning sites known from other opsins implying they are also unlikely to affect spectral tuning in anuran SWS2. The absorbance spectra of *O. pumilio* cones, however, does hint at the potential for substantial variation in anuran SWS2. This species, which was found to lack green rods, has cones with a λ_max_ of 466 nm which likely contain a red-shifted SWS2 pigment. Further studies are needed to explore the molecular mechanisms of this potential red-shift and other spectral tuning mechanisms in anuran SWS2 pigments.

### First Evidence of Visual Opsin Duplication in Amphibians

In the African Bullfrog, *Pyxicephalus adspersus*, we found the first evidence of a visual opsin gene duplication in amphibians with two copies of *LWS*, one on each of the two sex chromosomes. Visual opsin gene duplication is rare among tetrapods having previously only been reported in some marsupials where *RH1* was duplicated (Cowing et al. 2008) and in two primate lineages where *LWS* was duplicated (Carvalho et al. 2017), but is common in teleost fishes (Carleton et al. 2020). The location of the *LWS* duplicates on different sex chromosome in *P. adspersus* is distinct from the primate duplications where the two duplicates are found on the same sex chromosome (X), but could be functionally similar to the allelic variation in some primate LWS. In those primates, heterozygotes have two distinct *LWS* alleles on the X chromosomes that enable red-green colour discrimination in females, but not males (Carvalho et al. 2017). In *P. adspersus* the two *LWS* genes are on the Z and W chromosomes, respectively. Thus, males would have two copies of the same (Z) gene, while females would have two different copies (ZW) potentially enabling additional colour discrimination if the λ_max_ of the two genes has diverged. The Z and W *LWS* genes have several non-conservative changes at the 17 sites where they differ, but these are not at any known spectral tuning or positively selected sites. Thus, the potential impact of these changes on special tuning or other functional properties will require further study. A potential selective advantage of two *LWS* genes is also unclear, but could be related to a behaviour of females who will swim underwater to avoid smaller males in order to reach and mate with the larger, dominant male (Amphibiaweb 2021; ref). A second, red-shifted LWS pigment could provide an advantage in vision in the red-shifted freshwater environments, something that in the tadpoles of many species, and in a fully aquatic species such as *X. laevis*, is achieved through the use of the A2, instead of the A1, chromophore. This species is also one of a small number of diurnal frog species, which generally require further study to evaluate other potential adaptations to bright-light and colour vision in anurans.

### Potential Functional Adaptations for Dim-light vision in Anuran RH1 and SWS2

Most anurans are nocturnal, at least as adults, and thus we might expect their visual systems to be particularly adapted to vision in dim light conditions and at the morphological and cellular levels this appears to be the case. Many anurans have relatively large eyes (Thomas et al. 2020) as well as very large and numerous rod photoreceptors (Nilsson 1964). Additionally, most anurans have a second type of rod photoreceptor (SWS2 rods), which may further enhance visual sensitivity and enable colour discrimination at light levels where for most other animals only achromatic vision is possible (Yovanovich et al. 2017). We also identified several features at the molecular level that also may provide dim light adaptation. RH1 N83 has been identified as a dim light adaptation based on an accelerated formation of the active signaling state of the visual pigment. Mutations to N83 have also been shown to increase the time it takes for the chromophore to exit the binding pocket after light activation (retinal release rate), which could prolong the lifetime of the active state increasing light sensitivity (Bickelmann et al. 2012). All anurans in our sample had N83, including the two diurnal species in our dataset (*Pyxicephalus adspersus* and *Mantella baroni*), which could indicate this site has become fixed in frogs regardless of light environment. However, there is some disconnect between N83 and dim light environments because diurnal turtles and lizards have N83, while nocturnal crocodilians have D83 (Schott et al. 2018; RKS pers. obs.), which may indicate that there are more complex functional roles of substitutions at this site that require further study. A second site, 299, was also shown to affect retinal release rate where the substitutions S299A and A299S increased and decreased retinal release rates, respectively (Dungan and Chang 2017). Variation between S and A at site 299 also occurred in our sample of anurans, although interestingly S299 was not found in either of the diurnal species or those with partial daytime activity (e.g., *Lithobates* spp.). Thus species with the combination N83 and S299, which when mutated in bovine RH1 resulted in the slowest retinal release rate (Dungan and Chang 2017), were only found in nocturnal species and in particular included subfossorial and burrowing species (*Microhyla fissipes* and *Spea* and *Scaphiopus* spp.). Whether this is related to visual performance in these dim-light habitats remains to be tested.

We also found that all anurans in our sample had SWS2 with T47 regardless of activity pattern. This residue was shown to result in increased light sensitivity through increased dark state stability (i.e., low thermal isomerization rate) to levels nearly as high as RH1 (Kojima et al. 2017). Extremely high dark state stability of RH1 is one of the functional properties that enable single photon responses in rods (Lamb 2013), and thus is likely crucial for the function of SWS2 rods in dim light vision and would be necessary to achieve colour vision at scotopic light levels (Yovanovich et al. 2017). While it has not been explored, this increased sensitivity likely comes with a trade-off reducing response times and/or recovery rates which is much higher in cones (Lamb 2013). It is unknown whether all the anurans in our sample have SWS2 rods and the only species where SWS2 rods have been shown to be absent (*D. pumilio*; Siddiqi et al. 2004) lacks molecular data. Further studies will be needed to explore whether there is indeed a trade-off and if species lacking SWS2 rods have undergone a reversal at site 47.

## Conclusions

Anurans form a largely understudied but intriguing group of organisms for studies of visual system evolution, in part due to their reliance on visual cues and specialization for dim-light vision, including the unique use of two spectrally-distinct rod classes. Additionally, while most molecular vision studies have focused on organisms living in either aquatic or terrestrial light environments, anurans present an opportunity to study complex visual system metamorphosis between them. How potential tuning mechanisms such as amino acid variation at both known and potential tuning sites actually affects spectral sensitivity at the visual pigment, photoreceptor, and organism level requires further investigation. By contributing to our understanding of spectral tuning mechanisms in anuran visual systems, this study supports future investigative work into the fundamental questions of anuran visual ecology.

## Acknowledgments

We thank Don Pratt, Bea Clack, Rayna Bell, and Matthew Fujita for valuable feedback and assistance throughout this project. We also thank many SFA undergraduate and graduate students for assistance with collections and John Clare for providing the photograph of *Pyxicephalus adspersus*. Funding for this project was provided by Stephen F. Austin State University’s Office of Research and Sponsored Programs through the Faculty-Student Collaborative Research Program to LKP, MAK and JMG; the Society for Integrative and Comparative Biology through their Grants-in-Aid of Research program. RKS was funded through an NSF DEB award (DEB-#1655751). This work was completed under a Texas Parks and Wildlife Department Scientific Research Permit No. SPR-0118-004. The findings and conclusions in this article are those of the author(s) and do not necessarily represent the views of the U.S. Fish and Wildlife Service.

## Data Availability

Sequences generated with this study have been deposited to NCBI Genbank Accession #*TBD*. All other data associated with the study are available on Zenodo at http://doi.org/10.5281/zenodo.5252929

## Competing Interests

The authors declare that there are no competing interests.

## Appendix

### Appendix 1

Anuran opsin primers. The letters “F” and “R” at the end of each primer code indicate forward and reverse primers, respectively. Start and stop positions are based on complete *Xenopus laevis* reference sequences for each gene.

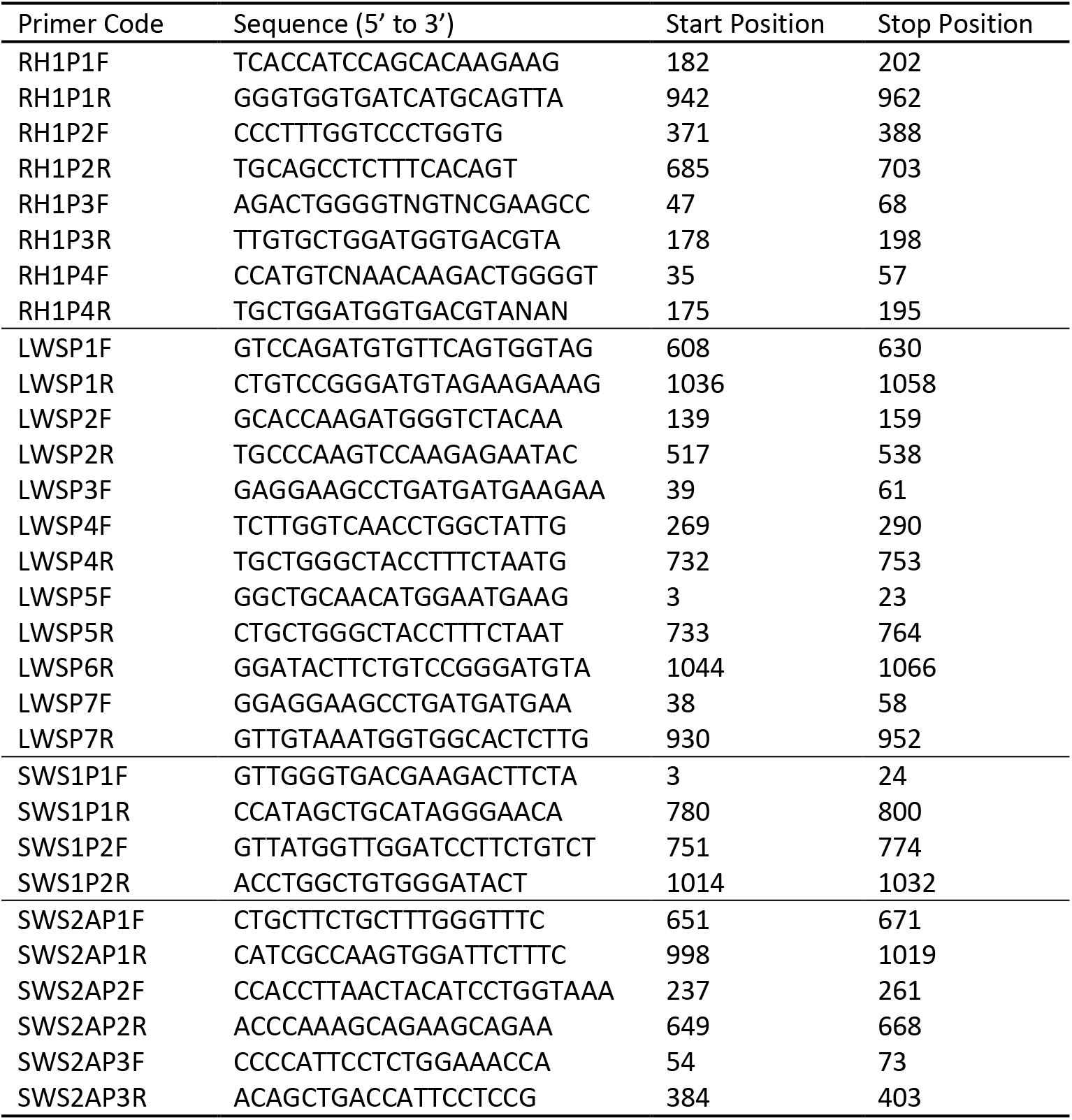

## Notes

### Competing Interest Statement

The authors have declared no competing interest.

